# Optimizing effect sizes and specificity trumps machine learning when building DNA methylation reference panels for cell-type deconvolution

**DOI:** 10.1101/2025.08.20.671293

**Authors:** Xiaolong Guo, Andrew E Teschendorff

## Abstract

Accurate cell-type deconvolution is critical for correct interpretation of Epigenome-Wide Association Studies. Such cell-type deconvolution involves estimating underlying cell-type fractions in a sample, which is accomplished using a DNA methylation reference panel built from sorted or single-cell DNAm data. Two competing approaches have emerged to build such reference panels, one which uses machine-learning, and another based on optimizing effect size and cell-type specificity. Here we demonstrate that the latter approach is preferable, because, owing to the relatively small number of sorted samples used in building panels, standard machine learning does not optimize effect size and cell-type specificity, causing the model to overfit and underperform when tested in independent data. Furthermore, adult blood panels built from cell-type specific hypomethylated markers improves inference when compared to panels built from hypermethylated ones. These insights provide important guidelines for optimizing the construction of future DNAm reference panels. To aid this task, we have added a function for building an optimized DNAm reference panel to our EpiDISH R-package.

## Introduction

The goal of Epigenome Wide Association Studies (EWAS) is to identify DNA methylation (DNAm) changes that are informative of disease risk factors and disease itself [1, 2]. Because DNAm is highly cell-type-specific [2, 3], measuring DNAm in bulk-tissue, as is most often done in epigenome studies, can cause statistical confounding [4]. This is because DNAm changes in bulk-tissue can arise from shifts in cell-type composition and not because of underlying DNAm changes in individual cell-types [3–6]. As shown by us and others [4, 7–9], this statistical challenge is best addressed using a cell-type deconvolution algorithm that first estimates the underlying cell-type proportions in a bulk-tissue sample, subsequently using these estimated fractions as covariates in linear models to identify differentially methylated cytosines (DMCs) that are not driven by changes in cell-type composition. The estimation of these cell-type fractions requires the construction of a DNAm reference panel, which contains representative DNAm profiles for all main underlying cell-types in the tissue of interest, and which is normally built by processing genome-wide DNAm profiles of sorted cell-types [9–12] or single-cells [13, 14]. Although studies have shown that the inference of cell-type fractions from such a DNAm reference panel is a relatively robust procedure [15], it is nevertheless important to ensure that DNAm reference panels are built in the most optimal way. This is particularly important in view of the growing urgent need for generating such panels for a wider range of different tissue-types at increased cellular resolution [14, 16–18]. In this regard, a key challenge is posed by the relatively small number of sorted samples available for each cell-type (typically less than 10) [10, 11], as generating larger numbers of sorted samples (say 25) for each of say 10-20 cell types, would require generation of several hundred samples. Similar limitations apply to single-cell DNAm (scDNAm) data, which is plagued by high sparsity and not yet widely available, with existing scDNAm-datasets only profiling samples from a small number of independent donors [13, 19, 20]. Thus, it is important to set up guidelines for the optimized construction of DNAm reference panels from what are typically small DNAm reference datasets.

Ideally, a DNAm reference panel should comprise DNAm markers that are (i) highly discriminatory between different cell types, (ii) stable within each cell type, and (iii) robust to variable factors such as age, race or genotype of the donors. As the deconvolution process essentially projects the methylation information of a bulk-tissue sample onto a feature space defined by the reference markers, if this space is poorly defined (i.e., markers lack true cell-type specificity or discriminatory power), the accuracy of the projection and estimation of cell-type fractions will be compromised. Traditionally, the selection of cell-type specific markers for a reference panel is based on identifying methylation sites with significant statistical differences between the various cell types. The first approach, due to Houseman et al. [9], independently fitted one-way analysis of variance (ANOVA) models to each CpG to test for differences in mean methylation levels across K purified cell types, selecting CpGs with the largest F-statistics to constitute the reference library/panel. However, the F-statistic prioritizes sites with large overall DNAm differences across all cell types, which can lead to the over-selection of CpGs that strongly differentiate one or a few cell types while neglecting others, resulting in imbalanced discriminatory power within the panel. To address the global limitation of the ANOVA method, subsequent studies shifted to using t-statistics (now called “Legacy method”, implemented in the Bioconductor package *minfi* using the function *pickCompProbes*) to select CpGs that are significantly hyper- or hypo-methylated in a specific cell type compared to all others [21]. Subsequent optimized feature selection frameworks have largely built upon the library generated by this “Legacy method” as their initial candidate pool of marker CpGs. Examples include iterative refinement strategies like IDOL and RESET [22, 23], as well as machine learning approaches such as Elastic Net and Random Forest [24]. However, marker CpGs selected purely on statistical significance ranking and machine-learning may potentially overlook the magnitude of the effect size, as a small effect can still be very informative if standard deviations are small. We reasoned that such small effect size CpGs are more likely to represent false positives, because based on the available evidence from large WGBS atlases [25], cell-type specific marker CpGs have large or reasonably large effect sizes. Moreover, selecting candidate marker CpGs using t-statistics does not guarantee maximum specificity for a given cell-type, specially if the number of sorted samples per cell-type varies significantly between cell-types.

To overcome these limitations, we recently introduced a novel marker selection strategy based on a joint t-statistic and “gap specificity score” (GSS) [26, 27]. This GSS metric is designed to directly prioritize the magnitude of the methylation difference—the ‘gap’—between a target cell type and all other cell types, thus maximizing effect size and specificity, while also ensuring that markers are selected based on statistical significance and hence a high within-cell-type stability. Here, we directly compare this GSS-based approach to the competing method described above based on using IDOL. We demonstrate that the GSS-based approach builds a more accurate and robust deconvolution reference panel. We further validate this approach by demonstrating its enhanced ability to detect known age-related changes in immune cells and to yield more biologically meaningful findings in epigenome-wide association studies (EWAS) of schizophrenia and type 1 diabetes. As such, the insights gained from our comparative analysis may serve as useful guidelines for the construction of future DNAm reference panels.

## Results

### Gap specificity score outperforms t-statistic by avoiding low effect size DMCs

The construction of a DNAm deconvolution reference matrix is detailed here, utilizing the blood reference library by Salas et al., which encompasses DNAm profiles from 12 sorted immune cell types measured using EPIC DNAm arrays [28] **(Methods, Fig.1a)**. Initially, for each CpG site, a two-sample t-test was performed to compare the mean DNAm of a given cell type against the collective remaining 11 cell types. For each cell type, CpG sites exhibiting a statistically significant difference, defined by a false discovery rate (FDR) less than 0.05, were identified as cell type-specific DMCs **(Fig.1a)**. These DMCs were then ranked for marker selection using one of two approaches. The first approach (the ‘legacy method’), selects ***N*** marker CpGs per cell type: the N/2 DMCs with the largest positive t-statistics and the N/2 DMCs with the largest negative t-statistics [21]. The second approach ranks DMCs by the gap specificity score (GSS). This score quantifies the methylation ‘gap’: for hypermethylated markers, it is the target cell type’s minimum methylation value minus the maximum methylation value taken over all other cell types; for hypomethylated markers, it is defined by the minimum methylation value over all other cell types minimum minus the target cell type’s maximum methylation value **(Fig.1a)**. Thus, in both cases, the GSS should ideally be positive and as large as possible, as this ensures specificity of the marker to the one given cell-type. GSS is designed to identify robust cell type-specific markers by prioritizing this ‘gap’ (reflecting effect size) alongside the marker’s cross-sample stability within the target cell type. For a fair comparison with the legacy method, we ranked hypermethylated and hypomethylated DMCs by GSS separately and selected the top N/2 from each category **(Fig.1a)**.

**Figure-1:**
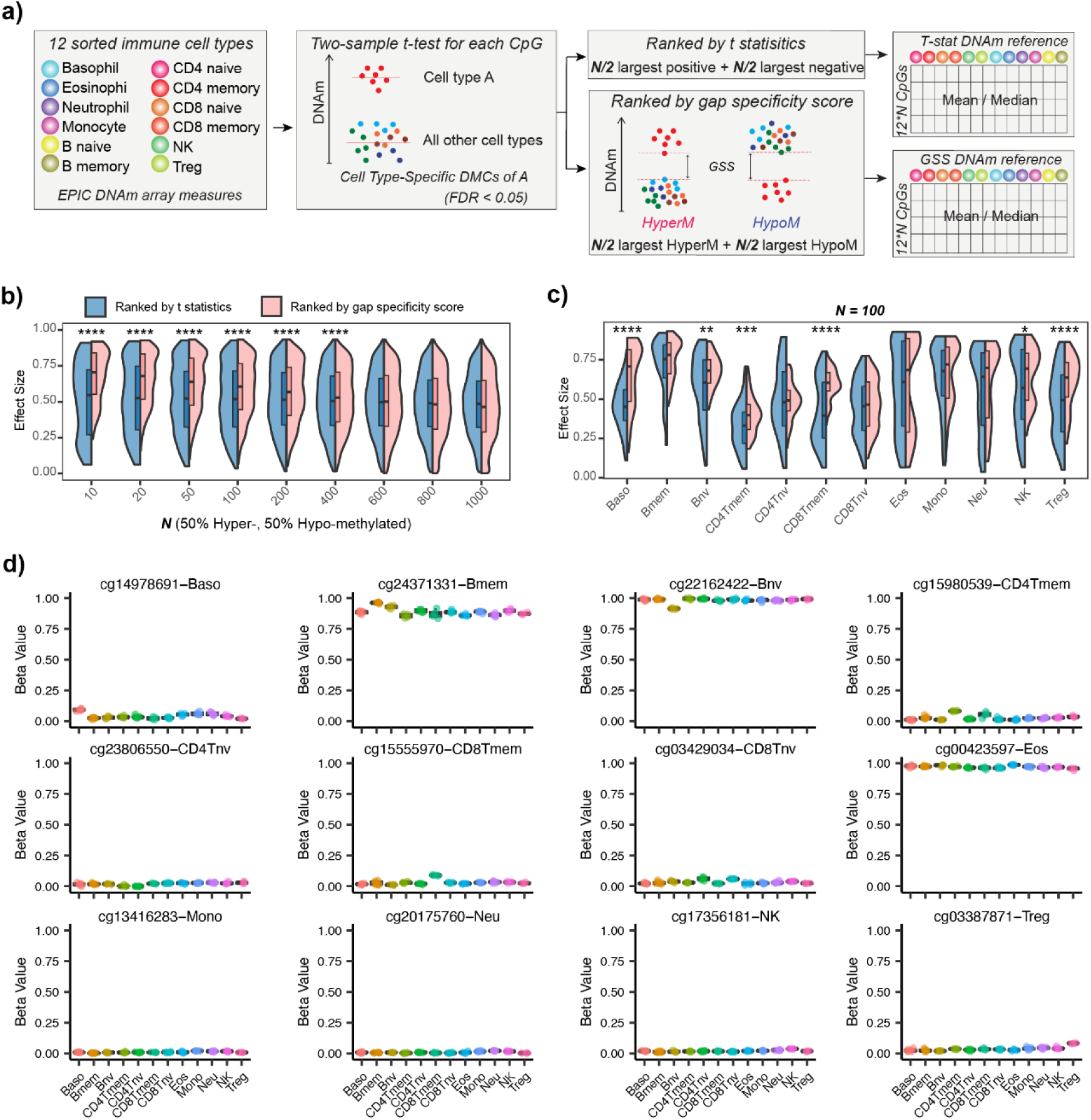
Gap specificity score avoids low-effect size DMCs obtained by ranking of t-statistics. **a)** Schematic diagram displaying the method of constructing deconvolution references using gap specificity score (GSS) and t-statistic. **b)** Violin plots display the distribution of effect sizes for DMCs ranked by gap specificity score versus t-statistic at each **N**. The x-axis, **N**, denotes the top N DMCs for each cell type, with N/2 hypermethylated and N/2 hypomethylated DMCs. The effect size for each DMC is calculated as the absolute mean methylation difference between the corresponding cell type and all other cell types. Statistical significance (****) indicates P < 1e-4, determined by a one-tailed Wilcoxon rank-sum test. **c)** Similar to b), directly compares the effect sizes of the top 100 DMCs obtained via gap specificity score ranking and t-statistic ranking for each cell type. Significance levels determined by a one-tailed Wilcoxon rank-sum test are indicated as: * P < 0.05, ** P < 0.01, *** P < 0.001, **** P < 1e-4. **d)** Boxplots show examples of low-effect-size markers obtained by the t-stat reference and that are retained after IDOL optimization, with one example presented for each cell type.

To determine if top DMCs ranked by the GSS exhibit a larger effect size compared to those ranked by t-statistics, we tested a series of incrementally increasing ***N*** values. The effect size of each DMC was defined as the absolute difference between the mean beta value of its target cell type and the mean beta value of the collective remaining 11 cell types. For ***N*** values ranging from 10 to 400, top DMCs ranked by the gap specificity score consistently exhibited significantly larger effect sizes than those ranked by the t-statistic **(Fig.1b)**. Next, we selected ***N***=100 to specifically compare the effect sizes from the two methods for each cell type. We observed that ranking based on t-statistics often included DMCs with notably low effect sizes, an issue especially prominent for lymphocyte subtypes **(Fig.1c**). Moreover, many of these low effect size DMCs find their way into DNAm reference panels built with machine-learning methods like IDOL (**Fig.1d**), thus highlighting a potential problem with using such methods to build DNAm reference panels.

### Low-effect size DMCs can negatively impact cell-type deconvolution accuracy

To confirm that these low effect size DMCs can negatively impact downstream cell-type deconvolution, we built DNAm reference matrices using the two different approaches (**Fig.1a**), subsequently applying them to deconvolve twelve experimental mixtures, each being a mixture of 12 immune-cell-types with known cell type proportions. We used EpiDISH [29] to estimate the underlying cell-type fractions, which were then compared to the true fractions by calculating the Pearson correlation coefficient (PCC) and root mean square error (RMSE) **(Methods, Fig.2a)**. While both ranking strategies yielded similar PCCs, the GSS-strategy consistently achieved a lower global RMSEs **(Fig.2a)**. Notably, the RMSE for memory CD4 T cells was markedly higher than those of other cell types (**Fig.2b**), consistent with their DMCs exhibiting the smallest effect sizes relative to other cell types **(Fig.1c)**.

**Figure-2:**
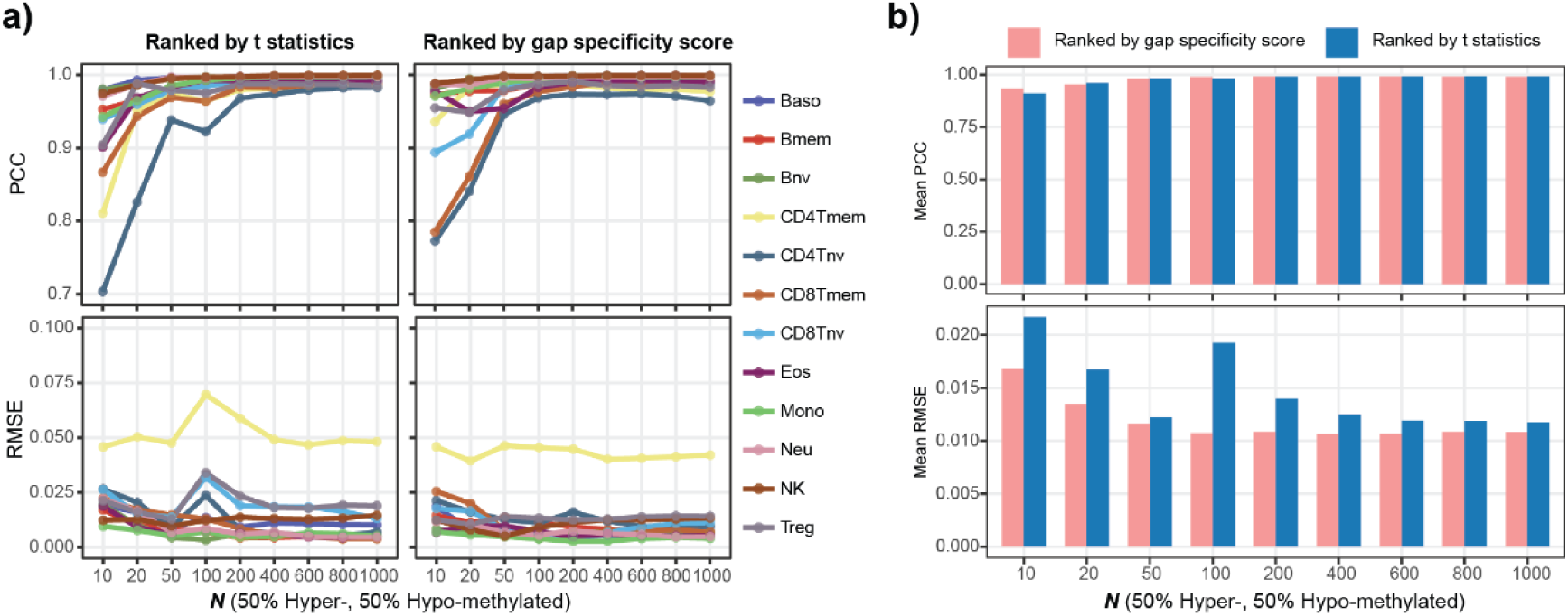
Low effect size DMCs negatively impact cell-type deconvolution accuracy. **a)** Line plots show the Pearson correlation coefficient (PCC) and root mean square error (RMSE) between true and estimated cell type proportions for 12 experimental mixtures at each **N**. **b)** Barplots summarize the average PCC and RMSE of a) across the 12 cell types at each **N**.

### Hypomethylated DMCs improve cell-type deconvolution accuracy

While the legacy method selected both hyper- and hypomethylated DMCs impartially, hypomethylated DMCs are more likely to be true cell-type-specific markers [25]. To further refine the GSS-based strategy, we thus asked if a reference constructed solely from top-ranked hypomethylated DMCs could outperform a reference combining both hyper- and hypomethylated DMCs, and a reference constructed using only top-ranked hypermethylated DMCs. Using a range of ***N*** values, the three distinct reference matrices were then compared in terms of their accuracy to deconvolve the twelve experimental 12-cell-type mixtures **(Fig. 3a)**. The reference constructed solely from hypermethylated DMCs yielded the lowest PCC values across all ***N*** values and the highest RMSE values when N > 100, compared to the other two reference types, indicating that only a small proportion of hypermethylated DMCs are effective for cell-type discrimination **(Fig. 3b)**. Conversely, using only top hypomethylated DMCs at N = 600 emerged as the best-performing reference, achieving the highest PCC and lowest RMSE **(Fig. 3b)**. Compared to the combined reference type, this ‘hypomethylated-only’ strategy resulted in a modest but clear reduction in RMSE, and notably improved deconvolution accuracy for memory CD4 T cells, which had previously exhibited poorer deconvolution accuracy **(Fig. 3a)**. While the ‘hypomethylated-only’ strategy performed best, the underlying combined reference type also achieved excellent performance (PCC ∼ 1, RMSE ∼ 0.01), demonstrating how the inclusion of many accurate markers can offset the presence of false positive low effect size markers **(Fig. 3b)**.

**Figure-3:**
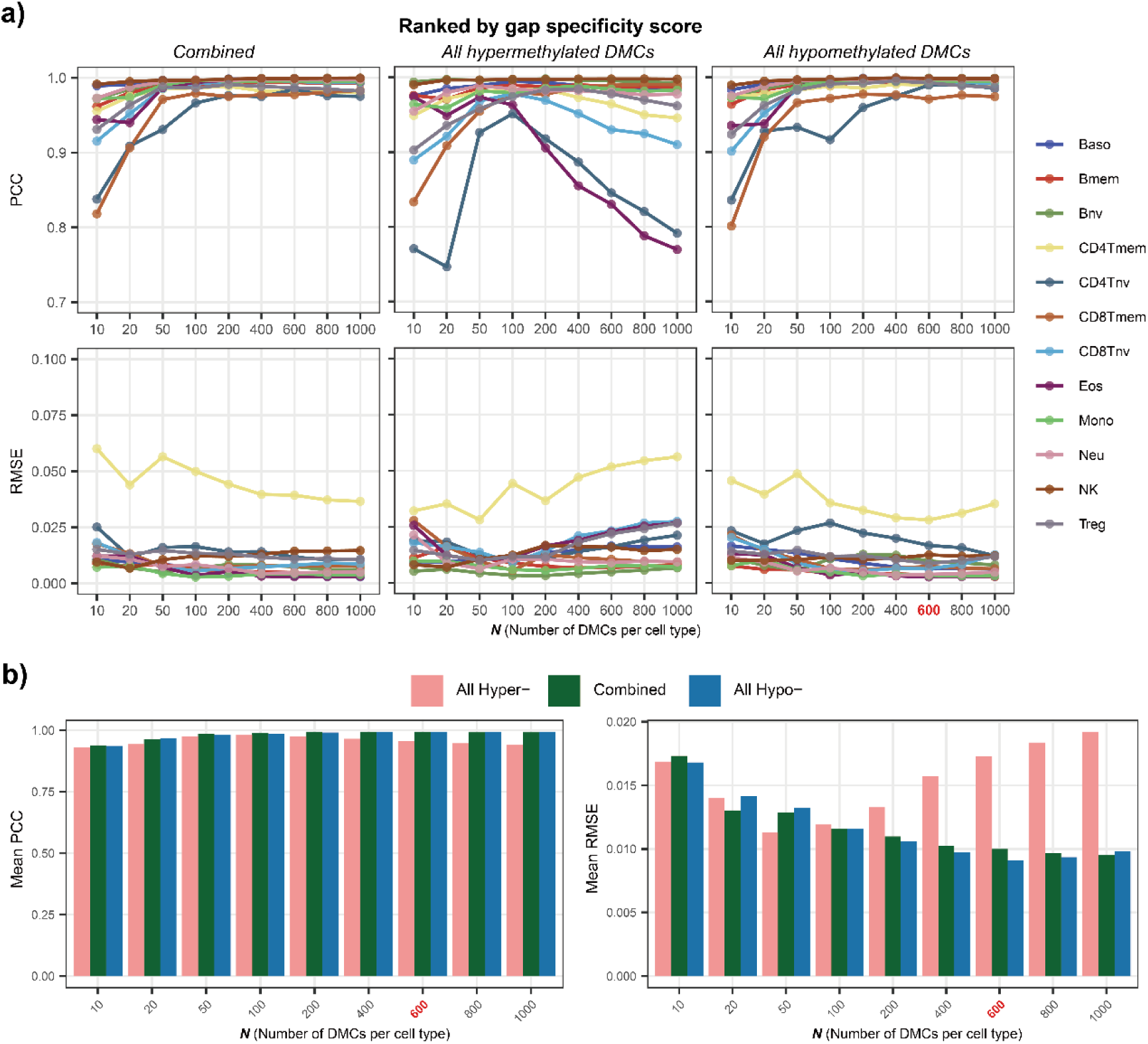
Exclusive use of hypomethylated markers improves deconvolution accuracy. **a)** Line plots display the Pearson correlation coefficient (PCC) and root mean square error (RMSE) of the true and estimated proportions for each cell type of 12 experimental mixtures. The “Combined” strategy refers to ranking solely based on the gap specificity score, without considering the direction of DMCs, to select the top DMCs for constructing the reference. The “All hypermethylated DMCs” and “All hypomethylated DMCs” strategies refer to ranking by gap specificity score and then using only hypermethylated DMCs or only hypomethylated DMCs, respectively, to build the reference. **b)** Barplots summarize the average PCC and RMSE for the 12 cell types at each library size as displayed in a). The position with the highest mean PCC and lowest RMSE has been marked in red on the x-axis.

### IDOL optimization for marker selection can overfit to small training data

Next, we sought to determine if the IDOL algorithm, which is often used to construct and optimize a DNAm reference panel [23, 28, 30, 31], could further improve the performance of a reference matrix built exclusively from top hypomethylated DMCs ranked by the GSS. Since both the legacy method and the GSS-strategy reached their best performance at N=600 **(Fig.2b, Fig.3b)**, we applied IDOL to the two reference matrices defined at this N-value, labeling them as ‘t-stat’ and ‘GSS-Hypo’, respectively. We note that IDOL also requires as input a training set comprising experimental mixtures generated from sorted immune cell types. Using the leave-one-out method, the probability of each CpG being selected is then updated based on whether the average R² and average RMSE of the estimated proportions for all cell types improves, continuing this process until convergence [23]. Thus, we applied the IDOL algorithm to optimize the t-stat and GSS-Hypo references, adopting the same training set and libSize = 1200 parameter as used by Salas et al.[28] **(Methods),** resulting in refined t-stat-IDOL and GSS-IDOL DNAm reference panels. Surprisingly, comparing the effect sizes of markers before and after IDOL optimization showed that IDOL has a tendency to select DMCs with lower effect sizes (**Fig.4a-b**), irrespective of whether we started out using the t-stat or GSS-Hypo reference panel. To explore this further, we assessed performance of the 4 reference matrices in the experimental mixtures from the training set as well as those of an independent validation set (**Methods**). Interestingly, the IDOL-optimized references demonstrated comparable PCCs in the training set but somewhat higher RMSEs (**Fig.4c**). Moreover, IDOL-optimized references performed worse in terms of both PCC and RMSE in the independent validation set (**Fig.4d**). Overall, ‘GSS-Hypo’ retained its superior deconvolution performance, even when compared to its IDOL-optimized version, indicating that IDOL may be overfitting slightly due to the relatively small number of experimental mixtures used in training.

**Figure-4:**
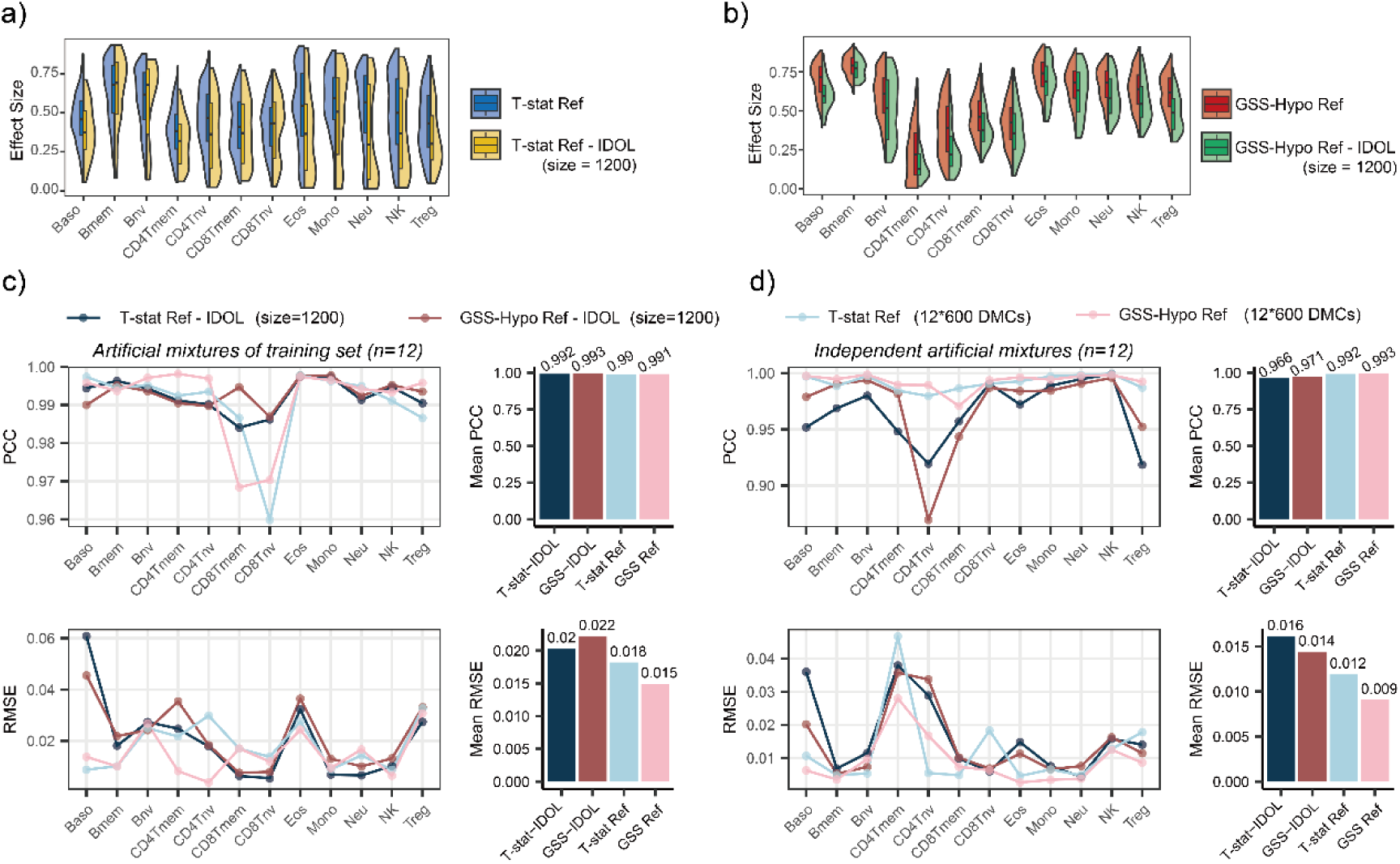
IDOL algorithm may not optimize DNAm reference panels. **a)** Violin plots displaying the distribution of effect sizes for DMCs from the ‘T-stat’ reference before and after IDOL optimization. **b)** as a), but for GSS-Hypo. **c)** Line plots illustrate the PCC and RMSE calculated between true and estimated proportions for each cell type across the 12 experimental mixtures in the training set. The adjacent bar plots summarize these results, displaying the average PCC and RMSE across all 12 cell types. **d)** as c), but for 12 independent experimental mixtures.

### Only the GSS-reference detects the known age-related increase of NK cells

As age increases, the immune cell composition of peripheral blood undergoes significant changes, characterized by shifts in the proportions of various subgroups, including the ratios of myeloid to lymphoid cells and memory to naive T cells [32–34]. To further compare the GSS-Hypo and t-stat reference panels, we examined their ability to capture known age-related changes in immune-cell type composition. Performing a meta-analysis across 15 adult blood cohorts, while adjusting for sex, we observed that for both reference panels, naive B-cell, CD8+ T-cell, and CD4+ T-cell fractions decreased with age, whilst corresponding memory T-cell fractions increased **(Fig.5a)**. These results align with findings from previous flow cytometry studies, particularly the pronounced decline of naive CD8⁺ T cells with age [32, 33]. However, only the GSS-Hypo panel identified a significant age-associated increase in the proportion of NK cells **(Fig.5a-b)**, consistent with previous reports [33, 35]. This demonstrates that the GSS-Hypo panel is more accurate in the estimation of cell-type fractions, as it can detect a known but subtle age-related change in NK proportions, whilst the t-stat panel fails to do so.

**Figure-5:**
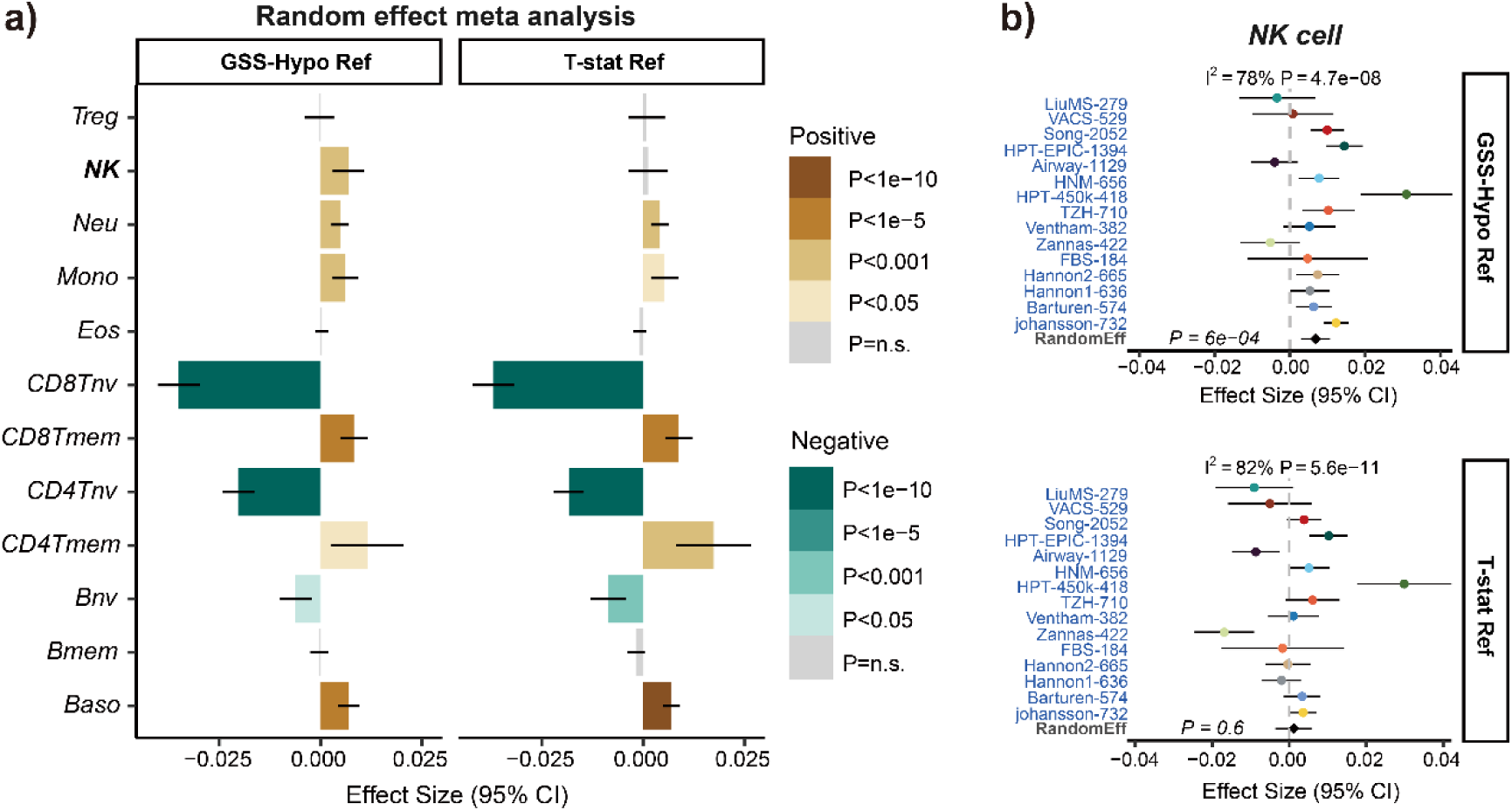
The t-stat reference panel does not detect the increased NK fraction with age. **a)** Barplots with 95% confidence intervals display the association of the 12 immune cell-type fractions estimated by the two references with chronological age, under a random effect model. **b)** Forest plots provide a more detailed illustration of the association of NK fractions with chronological age estimated by the two references in each cohort. P-value of a random effect meta-analysis model is given. Error bars represent 95% confidence intervals. Heterogeneity I^2^ indices and P-values are also given.

### GSS-reference panel improves interpretability of EWAS findings

Small differences in estimated cell-type fractions could also impact EWAS findings [29]. To test this, we performed an EWAS on two independent schizophrenia (SZ) case-control whole blood datasets from Hannon et al [36, 37], aiming to determine which reference matrix yielded more consistent disease-associated signals across cohorts **(Methods)**. In both the Hannon1 (discovery) and Hannon2 (validation) cohorts, we applied a multiple linear regression model, adjusting for covariates (e.g., age, sex) and cell-type fractions estimated by either the t-stat or GSS-Hypo reference. We identified significant SZ-associated DMCs in the Hannon1 cohort at a FDR < 0.05. These DMCs were then carried forward for validation in the Hannon2 cohort, with replication defined by a nominal P < 0.05. This two-stage approach yielded 322 and 264 replicated DMCs for the t-stat and GSS-Hypo references, respectively **(Fig. 6a)**. Although the GSS-panel resulted in fewer replicated associations, this does not imply more true positives. Moreover, although the two sets of replicated DMCs largely overlapped (n=217), we asked if the discordant DMCs could impact biological interpretation. To this end, we performed KEGG pathway enrichment analyses, separately on each set of DMCs, using missMethyl [38]. Analysis of the top ranked KEGG pathways revealed that the 264 SZ DMCs derived from the GSS-panel were more significantly enriched for known SZ-related pathways [39–45], compared to the 322 DMCs derived from the ‘t-stat’ reference, suggesting that the SZ-DMCs identified by the GSS-panel have a lower false positive rate **(Fig. 6b)**.

**Figure-6:**
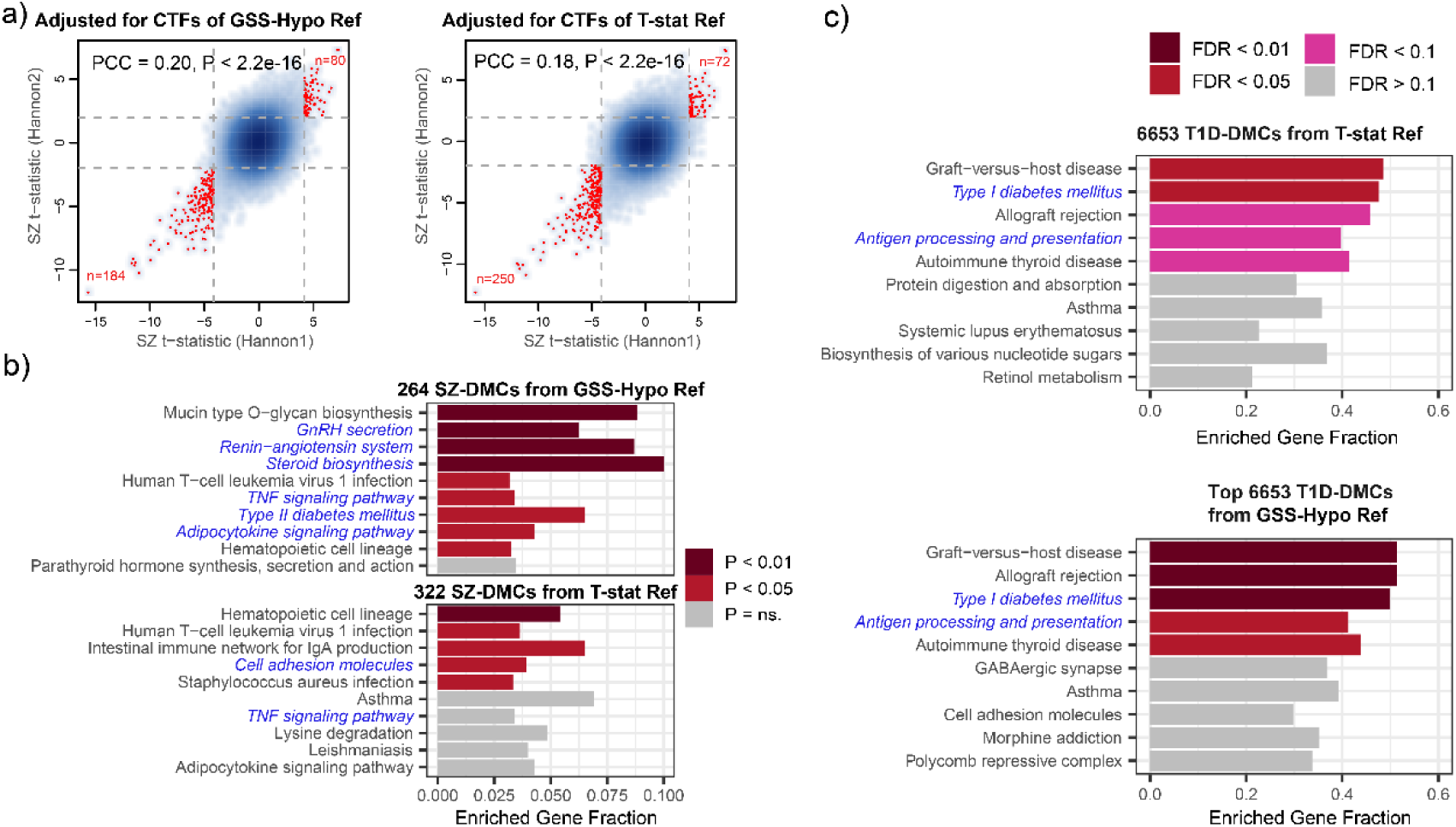
GSS-Hypo reference panel improves interpretability of EWAS hits. **a)** Scatter plots show the t-statistics of each CpG in EWAS conducted in two independent schizophrenia (SZ) case-control studies using fractions estimated with GSS-Hypo reference (left), T-stat reference (right). The horizontal gray dashed line represents the threshold where P < 0.05, while the vertical gray dashed line represents the threshold where FDR < 0.05. Red dots indicate points where significant in both studies. Pearson correlation coefficients and corresponding P values are also provided. **b)** Barplots depict the gene ratios for top KEGG pathways enriched among the replicated SZ-DMCs identified in a). **c)** Barplots depict the fraction of KEGG pathway genes that were found enriched among the T1D-DMCs. The number of T1D-DMCs in the bottom case has been matched to the number obtained using T-stat reference.

As a second validation, we analyzed a longitudinal DNAm dataset (Illumina 450k/EPIC) from a prospective study of type 1 diabetes (T1D) [46] **(Methods)**. This dataset, which has also been previously analyzed by us [27], comprises 395 blood samples collected from infants and children at multiple timepoints during the progression of islet autoimmunity (IA) prior to T1D diagnosis. Following a similar analytical approach as the SZ study, we used linear models to identify T1D-associated DMCs, adjusting for confounders including age, sex, timepoint, technical factors (chip and position), and the 12 immune cell fractions derived from either the GSS-Hypo or T-stat reference. The GSS-Hypo-adjusted analysis identified 9,589 T1D DMCs, compared to 6,653 DMCs identified using the t-stat reference (FDR < 0.05). We next assessed the biological relevance of these two DMC lists, restricting in both cases to the top 6653 DMCs to avoid any bias. Pathway enrichment analysis using missMethyl revealed that the top 6,653 DMCs derived from the GSS-panel exhibited more significant enrichment for the KEGG T1D pathway **(Fig. 6c)**. Overall, these data show that using the GSS-panel we obtain improvements in the enrichment of biologically relevant pathways.

## Discussion

Our study highlights the importance of a long-standing question in omic data science, namely, when selecting features do we rely on statistics only, or should we also take the magnitude of the effect into account? Generally speaking, using statistics and P-values to select features is of paramount importance. However, once features have been selected based on P-values, a less trivial question is whether significant features should be ranked by the statistic (or P-value) or whether they should be re-ranked according to effect size. The answer to this question largely depends on the biological problem and context. As demonstrated here, in the context of building DNAm reference panels for cell-type deconvolution, where one has to select features that are highly discriminatory of different cell-types, relying only on statistics and P-values to select and rank features is a suboptimal procedure, as it can inadvertently place low-effect size markers at the top of the ranked list, which, if subsequently not removed, can negatively impact downstream cell-type deconvolution. Indeed, our results clearly suggest that these low-effect size markers are likely false positives that arise because of the typically low number of sorted cell samples that are used when building DNAm reference panels. This is because of the high cost and difficulty of generating sorted samples for a relatively large number of donors and cell-types. Our GSS-strategy, which re-ranks features previously selected by statistics and P-values, by their cell-type specific effect size, can efficiently remove these false positives, improving the quality of the downstream cell-type deconvolution and statistical inference, as demonstrated here, not only in terms of the accuracy of the cell-type deconvolution itself, but also by improving the interpretability of EWAS findings.

Interestingly, we found that the IDOL algorithm did not optimize DNAm reference panels. When benchmarking the IDOL algorithm, Koestler et al. employed the *EstimateCellCounts* function from the *minfi* package, which relies on a reference constructed by the *pickCompProbes* function, which uses the t-statistic based legacy method for feature selection [21, 23]. This reference is limited to only 50 hypo-DMCs and 50 hyper-DMCs per cell type. As shown here, the legacy method underperforms with N=100 markers but improves as N rises to 600, where its deconvolution accuracy plateaus. On this basis, the application of IDOL optimization actually leads to a decline in performance, even when compared to the legacy method. According to Koestler’s description, increasing the algorithm’s search space (K*N, where K is the number of cell types) should theoretically improve performance [23]. On the other hand, it is important to emphasize that the reduced performance of the IDOL-derived reference panels was true irrespective of whether we re-ranked significant DMCs by the GSS or not, and that performance dropped further when testing them in independent datasets. Thus, a more likely explanation as to why IDOL may reduce performance is that it overfits to the experimental mixtures used in training. This is indeed highly plausible since the number of experimental mixtures (n=12) in the training set is small. Thus, we envisage that IDOL would be more valuable if it could be trained on a much larger set of experimental mixtures.

Another important insight from this work is that building DNAm reference panels exclusively using cell-type specific hypomethylated DMCs may further improve inference. This is consistent with the observation that the most cell-type specific CpGs are those that are unmethylated in the given cell-type, as indeed cell-type specific hypermethylated CpGs are relatively infrequent [25]. Thus, including hypermethylated markers may inadvertently increase the number of low-effect size false positive markers. However, we caution that this likely applies to only adult tissue-types and terminally differentiated cell-types [14, 25], because for other tissue-types like cord-blood, the presence of immature and progenitor cell-types may require hypermethylated markers to improve discrimination, as demonstrated recently by us [47]. Indeed, DNAm-atlases like that of Loyfer et al are focused on adult tissue and cell-types [25], which may lead to the misleading interpretation that most cell-type specific markers are always unmethylated, yet in early development DNAm undergoes substantial remodelling, the implication being that when comparing immature and progenitor/stem cell-types to each other, the amount of cell-type specific hypo and hypermethylation is much more comparable.

One potential caveat to our GSS-strategy is that we have not yet provided a concrete recommendation to the number of markers that should enter a DNAm reference panel. There are 4 important criteria that will determine what is a reasonable or optimal number of markers. First, it is desirable to have a balanced number of markers for each cell-type, as this ensures that the inference is not biased to the cell-types with more markers. A priori, when building a panel it is not known if this is even possible, and may require using different GSS thresholds per cell-type. Second, the number of markers per cell-type should be large enough so that one can reliably estimate fractions if say a fraction of these fail QC in independent datasets. Thus, redundancy in a panel is desirable as it confers robustness. On the other hand, using too many markers will eventually compromise their quality as measured by the GSS. Third, it is key that one maximizes the chance of the markers entering the panel be true positives, because if the proportion of high GSS true positive markers is high (say >70%), then there is no need for further optimization using an algorithm such as IDOL. Indeed, it is worth remembering that the purpose of optimization through cross-validation is to remove false positives. If however the original selection of markers is characterized by high GSS-values (say >0.5), then these are extremely unlikely to be false positives, in which case any optimization may only be counterproductive, specially if the training or optimization is performed on small datasets. Indeed, potential confounders like age, smoking, genotype or ethnicity generally only account for much smaller effect sizes [47, 48]. To make this point clear, a method like IDOL would be necessary only if the original marker selection contains a significant number of false positives driven by such confounders, yet as shown here, IDOL would also require a larger training set. Fourth, the number of markers per cell-type is also a function of the number of cell-types in the panel. The more cell-types we have, the more markers we may need, since distinguishing more similar cell-types from each other (say classical from non-classical monocytes) may require a larger number of lower GSS features, if say high GSS features discriminating these similar cell-types do not exist. In summary, the number of markers per cell-type is a complex function of many factors, and not necessarily amenable to an optimization procedure, specially if the number of training set samples is small. However, based on our experience building and successfully validating DNAm reference panels [7, 15, 29, 47, 49], we advise on the order of around 50 GSS > 0.3 markers per cell-type, so ∼600 markers for a 12 cell-type panel.

Although our GSS-strategy led to improvements in statistical inference in the context of 3 different EWAS analyses, we acknowledge that performance of the various DNAm reference panels is quite similar. This is consistent with the underlying robustness of the cell-type deconvolution estimation procedure [15]: the DNAm reference panel can tolerate up to 30% errors/false positives without a significant degradation in performance. Nevertheless, as future efforts will need to generate DNAm reference matrices for larger numbers of cell-types [18] using say sorted or single-cell DNAm data [14, 19, 20], this will require accurate deconvolution for an increasing number of cell-types, which will place more stringent demands on the reliability and quality of a DNAm reference panel. Thus, the insights gained here can provide guidelines to help build future and more accurate DNAm reference panels. To aid the community, we now provide within our EpiDISH R-package [29], an R-function to build DNAm reference panels from sorted or single-cell DNAm data using our GSS-strategy.

In summary, sole reliance on statistics and P-values to select features for building DNAm reference panels leads to suboptimal cell-type deconvolution, even if followed-up with a machine-learning optimization strategy. This is because of the inclusion of low effect size markers, as well as overfitting to the inevitably small datasets comprising independent sorted or single-cell DNAm data. As shown here, the GSS-strategy to select a reasonable number of markers with high specificity and large effect size optimizes cell-type deconvolution tasks.

## Methods

### Sorted immune cell types and artificial mixtures

*Salas et al.* (GSE167998)[28]: This dataset comprises EPIC profiles of 12 sorted immune-cell subsets, totaling 56 sorted samples: 6 basophils (Baso), 6 memory B-cells (Bmem), 4 naive B-cells (Bnv), 4 memory CD4+ T-cells (CD4Tmem), 5 naive CD4+ T-cells (CD4Tnv), 4 memory CD8+ T-cells (CD8Tmem), 5 naive CD8+ T-cells (CD8Tnv), 4 eosinophils (Eos), 5 monocytes (Mono), 6 neutrophils (Neu), 4 natural killer (NK) cells, and 3 regulatory T-cells (Treg). Additionally, there are 12 artificial mixture samples composed of these 12 cell types, with known mixing proportions for each cell type. The dataset had been previously processed by Luo et al.[50]

*Salas et al.* (GSE182379)[28]: This dataset comprises EPIC profiles of 12 artificial mixture samples of 12 immune cell types, with known mixing proportions for each cell type. Consistent with the data processing pipeline used by Luo et al. for the GSE167998 dataset: The idat files were downloaded and processed using the minfi R-package. We retained 775,940 probes with significantly detected values across all 12 samples. The resulting beta-valued data matrix was then adjusted for type-2 probe bias using BMIQ. These two artificial mixture datasets all come from the cell mixture reconstruction experiment. Specifically, DNA extracted from purified 12 immune cell subtypes were mixed in preplanned proportions to reconstruct blood samples.

### Cell Mixture Deconvolution

Previous studies have clearly described the principles of Cell Mixture Deconvolution, which will not be elaborated on here [22, 28, 29, 51]. Ridder et al. have demonstrated that the RPC method of EpiDISH is currently the best-performing algorithm among 16 deconvolution algorithms[52]. Therefore, we estimated corresponding cell-type fractions using the EpiDISH Bioconductor R-package[29, 53]. Specifically, we ran the epidish function with “RPC” as the method and maxit = 500.

### IDOL algorithm

Koestler et al. have provided the source code for the core IDOL function, IDOLoptimize, and the IDOL R package in their GitHub repository https://github.com/immunomethylomics/IDOL [23]. As performed by Salas et al.[28], we constructed different DMC libraries using the sorted immune cell types from GSE167998, then optimized them using the artificial mixtures from GSE167998, and subsequently conducted independent validation using the artificial mixtures from GSE182379. The library size parameter, libSize = 1200, was determined as the optimal size by Salas et al. based on testing ranging from 250 to 3000 CpGs[28].

### Human adult whole blood DNA methylation datasets

All the cohorts used here have previously been normalized and analyzed by us [49], including: (1) *LiuMS (GSE106648)*: 279 HM450k peripheral blood (PB) samples, with age range from 16 to 66 (140 multiple sclerosis patients + 139 controls) [54]; (2) *Song (GSE169156)*, 2052 normal EPIC PB samples, with age range from 18 to 66 [55]; (3) *HPT-EPIC & HPT-450k (GSE210255 & GSE210254)*: 1394 normal PB samples (EPIC set, age range from 21 to 87) and 418 normal PB samples (450k set, age range from 34 to 91) [56]; (4) *Barturen (GSE179325)*: 574 normal EPIC PB samples, with age range from 19 to 103 [57]; (5) *Airwave (GSE147740)*: 1129 normal EPIC PB samples, with age range from 25 to 60 [58]; (6) *VACS (GSE117860)*: 529 HM450k samples from whole blood in HIV-positive men, with age range from 25 to 75 [59]; (7) *Ventham (GSE87648)*: HM450k PB samples from 204 newly-diagnosed IBD cases and 178 controls, with age range from 17 to 79, and 2 samples have no age information [60]; (8) *Hannon-1 and 2 (GSE80417 & GSE84727):* 675 HM450k PB samples with age range from 18 to 90 (353 schizophrenia cases and 322 controls), and 847 HM450k PB samples with age range from 18 to 81 (414 schizophrenia cases and 433 controls) [36, 37]; (9) *Zannas (GSE72680)*: 422 normal HM450k PB samples, with age range from 18 to 77 [61]; (10) *Flanagan/FBS (GSE61151)*: 184 normal HM450k PB samples, with age range from 35 to 83 [62]; (11) *Johansson (GSE87571)*: 732 normal HM450k PB samples, with age range from 14 to 94, and 3 samples have no age information [63]; (12) *TZH (OEP000260)*: 710 normal EPIC PB samples, with age range from 19 to 71 [64]; (13) *HNM (GSE40279)*: 656 normal HM450k PB samples, with age range from 19 to 101 [65].

### DNAm datasets of type-1 diabetes

DAISY study[46]: The Illumina DNAm dataset was downloaded from the GEO data repository (GEO accession GSE142512). This dataset contains 395 samples from 174 individuals. The ages of the individuals range from 6 months to 22 years, with the majority being under 10 years old. A total of 211 samples were sequenced using the EPIC array, and 184 samples were sequenced using the 450k array. Half of the individuals were eventually diagnosed with type 1 diabetes, but all samples were taken before diagnosis.

### Meta-analyses

In each cohort, we standardized the fractions of each cell type to unit variance. We then assessed associations between the standardized cell type fractions and chronological age using a multivariate linear regression. For specific cohorts where additional covariates were available such as gender, disease and smoking status, the multivariate regression models include these factors as covariates. The covariate information used for each cohort is described in detail in Luo et al [49]. For each regression, we extracted the corresponding effect size, standard error, Student’s *t* test, and *P*-value. We then performed a fixed and random effect inverse variance meta-analysis using the metagen function implemented in the meta R-package (version 7.0-0)[66].

## Data Availability

The following DNAm datasets analyzed here are publicly available from the NCBI GEO website https://www.ncbi.nlm.nih.gov/geo/ under the accession numbers GSE40279 (HNM), GSE106648 (LiuMS), GSE169156 (Song), GSE210255 (HPT-EPIC), GSE210254 (HPT-450k), GSE179325 (Barturen), GSE147740 (Airwave), GSE117860 (VACS), GSE87648 (Ventham), GSE84727 (Hannon2), GSE80417 (Hannon1), GSE72680 (Zannas), GSE61151 (Flanagan/FBS), GSE87571 (Johansson), GSE142512 (DAISY study), GSE167998 (Salas), GSE182379 (Salas), The Illumina EPIC DNAm data for the TZH cohort can be viewed at NODE under accession number OEP000260, or directly at (https://www.biosino.org/node/project/detail/OEP000260), and accessed by submitting a request for data-access.

## Code Availability

A function *ConstructDNAmPanel.R* has been added to our EpiDISH R-package freely available from https://github.com/sjczheng/EpiDISH.

## Acknowledgements

This project was funded by the National Natural Science Foundation of China grants 32170652 and 32370699, as well as a Research Fellowship for International Scientists (RFIS, W2431024).

## Author contributions

Study was conceived by AET. Statistical and bioinformatic analyses were performed by XG. Manuscript was written by XG and AET.

## Ethics approval

Not applicable as we only analyzed publicly available data.

## References

1. Rakyan VK, Down TA, Balding DJ, Beck S: Epigenome-wide association studies for common human diseases. Nat Rev Genet 2011, 12:529–541.

2. Lappalainen T, Greally JM: Associating cellular epigenetic models with human phenotypes. Nat Rev Genet 2017, 18:441–451.

3. Teschendorff AE, Relton CL: Statistical and integrative system-level analysis of DNA methylation data. Nat Rev Genet 2018, 19:129–147.

4. Liu Y, Aryee MJ, Padyukov L, Fallin MD, Hesselberg E, Runarsson A, Reinius L, Acevedo N, Taub M, Ronninger M, et al: Epigenome-wide association data implicate DNA methylation as an intermediary of genetic risk in rheumatoid arthritis. Nat Biotechnol 2013, 31:142–147.

5. Jaffe AE, Irizarry RA: Accounting for cellular heterogeneity is critical in epigenome-wide association studies. Genome Biol 2014, 15:R31.

6. Zhang Z, Christensen BC, Salas LA: Recalibrate concepts of epigenetic aging clocks in human health. Aging (Albany NY) 2024, 16:11125–11127.

7. Zheng SC, Beck S, Jaffe AE, Koestler DC, Hansen KD, Houseman AE, Irizarry RA, Teschendorff AE: Correcting for cell-type heterogeneity in epigenome-wide association studies: revisiting previous analyses. Nat Methods 2017, 14:216–217.

8. Teschendorff AE, Zheng SC: Cell-type deconvolution in epigenome-wide association studies: a review and recommendations. Epigenomics 2017, 9:757–768.

9. Houseman EA, Accomando WP, Koestler DC, Christensen BC, Marsit CJ, Nelson HH, Wiencke JK, Kelsey KT: DNA methylation arrays as surrogate measures of cell mixture distribution. BMC Bioinformatics 2012, 13:86.

10. Salas LA, Zhang Z, Koestler DC, Butler RA, Hansen HM, Molinaro AM, Wiencke JK, Kelsey KT, Christensen BC: Enhanced cell deconvolution of peripheral blood using DNA methylation for high-resolution immune profiling. Nat Commun 2022, 13:761.

11. Gervin K, Salas LA, Bakulski KM, van Zelm MC, Koestler DC, Wiencke JK, Duijts L, Moll HA, Kelsey KT, Kobor MS, et al: Systematic evaluation and validation of reference and library selection methods for deconvolution of cord blood DNA methylation data. Clin Epigenetics 2019, 11:125.

12. Salas LA, Koestler DC, Butler RA, Hansen HM, Wiencke JK, Kelsey KT, Christensen BC: An optimized library for reference-based deconvolution of whole-blood biospecimens assayed using the Illumina HumanMethylationEPIC BeadArray. Genome Biol 2018, 19:64.

13. Cai M, Zhou J, McKennan C, Wang J: scMD facilitates cell type deconvolution using single-cell DNA methylation references. Commun Biol 2024, 7:1.

14. Zhu T, Liu J, Beck S, Pan S, Capper D, Lechner M, Thirlwell C, Breeze CE, Teschendorff AE: A pan-tissue DNA methylation atlas enables in silico decomposition of human tissue methylomes at cell-type resolution. Nat Methods 2022, 19:296–306.

15. Zheng SC, Webster AP, Dong D, Feber A, Graham DG, Sullivan R, Jevons S, Lovat LB, Beck S, Widschwendter M, Teschendorff AE: A novel cell-type deconvolution algorithm reveals substantial contamination by immune cells in saliva, buccal and cervix. Epigenomics 2018, 10:925–940.

16. Zhang Z, Lu Y, Vosoughi S, Levy JJ, Christensen BC, Salas LA: HiTAIC: hierarchical tumor artificial intelligence classifier traces tissue of origin and tumor type in primary and metastasized tumors using DNA methylation. NAR Cancer 2023, 5:zcad017.

17. Zhang Z, Wiencke JK, Kelsey KT, Koestler DC, Christensen BC, Salas LA: HiTIMED: hierarchical tumor immune microenvironment epigenetic deconvolution for accurate cell type resolution in the tumor microenvironment using tumor-type-specific DNA methylation data. J Transl Med 2022, 20:516.

18. Pike SC, Wiencke JK, Zhang Z, Molinaro AM, Hansen HM, Koestler DC, Christensen BC, Kelsey KT, Salas LA: Glioma immune microenvironment composition calculator (GIMiCC): a method of estimating the proportions of eighteen cell types from DNA methylation microarray data. Acta Neuropathol Commun 2024, 12:170.

19. Chien JF, Liu H, Wang BA, Luo C, Bartlett A, Castanon R, Johnson ND, Nery JR, Osteen J, Li J, et al: Cell-type-specific effects of age and sex on human cortical neurons. Neuron 2024, 112:2524–2539 e2525.

20. Lee DS, Luo C, Zhou J, Chandran S, Rivkin A, Bartlett A, Nery JR, Fitzpatrick C, O’Connor C, Dixon JR, Ecker JR: Simultaneous profiling of 3D genome structure and DNA methylation in single human cells. Nat Methods 2019, 16:999–1006.

21. Aryee MJ, Jaffe AE, Corrada-Bravo H, Ladd-Acosta C, Feinberg AP, Hansen KD, Irizarry RA: Minfi: a flexible and comprehensive Bioconductor package for the analysis of Infinium DNA methylation microarrays. Bioinformatics 2014, 30:1363–1369.

22. Bell-Glenn S, Thompson JA, Salas LA, Koestler DC: A Novel Framework for the Identification of Reference DNA Methylation Libraries for Reference-Based Deconvolution of Cellular Mixtures. Frontiers in Bioinformatics 2022, Volume 2 - 2022.

23. Koestler DC, Jones MJ, Usset J, Christensen BC, Butler RA, Kobor MS, Wiencke JK, Kelsey KT: Improving cell mixture deconvolution by identifying optimal DNA methylation libraries (IDOL). BMC Bioinformatics 2016, 17:120.

24. Arneson D, Yang X, Wang K: MethylResolver—a method for deconvoluting bulk DNA methylation profiles into known and unknown cell contents. Communications Biology 2020, 3:422.

25. Loyfer N, Magenheim J, Peretz A, Cann G, Bredno J, Klochendler A, Fox-Fisher I, Shabi-Porat S, Hecht M, Pelet T, et al: A DNA methylation atlas of normal human cell types. Nature 2023, 613:355–364.

26. Teschendorff AE, Zhu T, Breeze CE, Beck S: EPISCORE: cell type deconvolution of bulk tissue DNA methylomes from single-cell RNA-Seq data. Genome Biol 2020, 21:221.

27. Guo X, Sulaiman M, Neumann A, Zheng SC, Cecil CAM, Teschendorff AE, Heijmans BT: Unified high-resolution immune cell fraction estimation in blood tissue from birth to old age. Genome Medicine 2025, 17:63.

28. Salas LA, Zhang Z, Koestler DC, Butler RA, Hansen HM, Molinaro AM, Wiencke JK, Kelsey KT, Christensen BC: Enhanced cell deconvolution of peripheral blood using DNA methylation for high-resolution immune profiling. Nature Communications 2022, 13:761.

29. Teschendorff AE, Breeze CE, Zheng SC, Beck S: A comparison of reference-based algorithms for correcting cell-type heterogeneity in Epigenome-Wide Association Studies. BMC Bioinformatics 2017, 18:105.

30. Gervin K, Salas LA, Bakulski KM, van Zelm MC, Koestler DC, Wiencke JK, Duijts L, Moll HA, Kelsey KT, Kobor MS, et al: Systematic evaluation and validation of reference and library selection methods for deconvolution of cord blood DNA methylation data. Clinical Epigenetics 2019, 11:125.

31. Salas LA, Koestler DC, Butler RA, Hansen HM, Wiencke JK, Kelsey KT, Christensen BC: An optimized library for reference-based deconvolution of whole-blood biospecimens assayed using the Illumina HumanMethylationEPIC BeadArray. Genome Biology 2018, 19:64.

32. Li M, Yao D, Zeng X, Kasakovski D, Zhang Y, Chen S, Zha X, Li Y, Xu L: Age related human T cell subset evolution and senescence. Immunity & Ageing 2019, 16:24.

33. Lin Y, Kim J, Metter EJ, Nguyen H, Truong T, Lustig A, Ferrucci L, Weng N-p: Changes in blood lymphocyte numbers with age in vivo and their association with the levels of cytokines/cytokine receptors. Immunity & Ageing 2016, 13:24.

34. Zhang L, Mack R, Breslin P, Zhang J: Molecular and cellular mechanisms of aging in hematopoietic stem cells and their niches. Journal of Hematology & Oncology 2020, 13:157.

35. Brauning A, Rae M, Zhu G, Fulton E, Admasu TD, Stolzing A, Sharma A: Aging of the Immune System: Focus on Natural Killer Cells Phenotype and Functions. Cells 2022, 11:1017.

36. Hannon E, Dempster E, Viana J, Burrage J, Smith AR, Macdonald R, St Clair D, Mustard C, Breen G, Therman S, et al: An integrated genetic-epigenetic analysis of schizophrenia: evidence for co-localization of genetic associations and differential DNA methylation. Genome Biol 2016, 17:176.

37. Hannon E, Dempster EL, Mansell G, Burrage J, Bass N, Bohlken MM, Corvin A, Curtis CJ, Dempster D, Di Forti M, et al: DNA methylation meta-analysis reveals cellular alterations in psychosis and markers of treatment-resistant schizophrenia. Elife 2021, 10.

38. Phipson B, Maksimovic J, Oshlack A: missMethyl: an R package for analyzing data from Illumina’s HumanMethylation450 platform. Bioinformatics 2016, 32:286–288.

39. Sheikh MA, O’Connell KS, Lekva T, Szabo A, Akkouh IA, Osete JR, Agartz I, Engh JA, Andreou D, Boye B, et al: Systemic Cell Adhesion Molecules in Severe Mental Illness: Potential Role of Intercellular CAM-1 in Linking Peripheral and Neuroinflammation. Biological Psychiatry 2023, 93:187–196.

40. Hoseth EZ, Ueland T, Dieset I, Birnbaum R, Shin JH, Kleinman JE, Hyde TM, Mørch RH, Hope S, Lekva T, et al: A Study of TNF Pathway Activation in Schizophrenia and Bipolar Disorder in Plasma and Brain Tissue. Schizophrenia Bulletin 2017, 43:881–890.

41. Dong K, Wang S, Qu C, Zheng K, Sun P: Schizophrenia and type 2 diabetes risk: a systematic review and meta-analysis. Frontiers in Endocrinology 2024, Volume 15 - 2024.

42. Ferrier IN, Cotes PM, Crow TJ, Johnstone EC: Gonadotrophin secretion abnormalities in chronic schizophrenia. Psychological Medicine 1982, 12:263–273.

43. Yasuno F, Suhara T, Ichimiya T, Takano A, Ando T, Okubo Y: Decreased 5-HT1A receptor binding in amygdala of schizophrenia. Biological Psychiatry 2004, 55:439–444.

44. Gadelha A, Vendramini AM, Yonamine CM, Nering M, Berberian A, Suiama MA, Oliveira V, Lima-Landman MT, Breen G, Bressan RA, Abílio V, Hayashi MAF: Convergent evidences from human and animal studies implicate angiotensin I-converting enzyme activity in cognitive performance in schizophrenia. Translational Psychiatry 2015, 5:e691–e691.

45. Mednova IA, Boiko AS, Kornetova EG, Parshukova DA, Semke AV, Bokhan NA, Loonen AJM, Ivanova SA: Adipocytokines and Metabolic Syndrome in Patients with Schizophrenia. In Metabolites, vol. 10; 2020.

46. Johnson RK, Vanderlinden LA, Dong F, Carry PM, Seifert J, Waugh K, Shorrosh H, Fingerlin T, Frohnert BI, Yang IV, et al: Longitudinal DNA methylation differences precede type 1 diabetes. Scientific Reports 2020, 10:3721.

47. Guo X, Sulaiman M, Neumann A, Zheng SC, Cecil CAM, Teschendorff AE, Heijmans BT: Unified high-resolution immune cell fraction estimation in blood tissue from birth to old age. Genome Med 2025, 17:63.

48. Peng Q, Liu X, Li W, Jing H, Li J, Gao X, Luo Q, Breeze CE, Pan S, Zheng Q, et al: Analysis of blood methylation quantitative trait loci in East Asians reveals ancestry-specific impacts on complex traits. Nat Genet 2024, 56:846–860.

49. Luo Q, Dwaraka VB, Chen Q, Tong H, Zhu T, Seale K, Raffaele JM, Zheng SC, Mendez TL, Chen Y, et al: A meta-analysis of immune-cell fractions at high resolution reveals novel associations with common phenotypes and health outcomes. Genome Med 2023, 15:59.

50. Luo Q, Dwaraka VB, Chen Q, Tong H, Zhu T, Seale K, Raffaele JM, Zheng SC, Mendez TL, Chen Y, et al: A meta-analysis of immune-cell fractions at high resolution reveals novel associations with common phenotypes and health outcomes. Genome Medicine 2023, 15:59.

51. Houseman EA, Accomando WP, Koestler DC, Christensen BC, Marsit CJ, Nelson HH, Wiencke JK, Kelsey KT: DNA methylation arrays as surrogate measures of cell mixture distribution. BMC Bioinformatics 2012, 13:86.

52. De Ridder K, Che H, Leroy K, Thienpont B: Benchmarking of methods for DNA methylome deconvolution. Nature Communications 2024, 15:4134.

53. Zheng SC, Breeze CE, Beck S, Dong D, Zhu T, Ma L, Ye W, Zhang G, Teschendorff AE: EpiDISH web server: Epigenetic Dissection of Intra-Sample-Heterogeneity with online GUI. Bioinformatics 2020, 36:1950–1951.

54. Kular L, Liu Y, Ruhrmann S, Zheleznyakova G, Marabita F, Gomez-Cabrero D, James T, Ewing E, Linden M, Gornikiewicz B, et al: DNA methylation as a mediator of HLA-DRB1*15:01 and a protective variant in multiple sclerosis. Nat Commun 2018, 9:2397.

55. Song N, Hsu CW, Pan H, Zheng Y, Hou L, Sim JA, Li Z, Mulder H, Easton J, Walker E, et al: Persistent variations of blood DNA methylation associated with treatment exposures and risk for cardiometabolic outcomes in long-term survivors of childhood cancer in the St. Jude Lifetime Cohort. Genome Med 2021, 13:53.

56. Shang L, Zhao W, Wang YZ, Li Z, Choi JJ, Kho M, Mosley TH, Kardia SLR, Smith JA, Zhou X: meQTL mapping in the GENOA study reveals genetic determinants of DNA methylation in African Americans. Nat Commun 2023, 14:2711.

57. Barturen G, Carnero-Montoro E, Martinez-Bueno M, Rojo-Rello S, Sobrino B, Porras-Perales O, Alcantara-Dominguez C, Bernardo D, Alarcon-Riquelme ME: Whole blood DNA methylation analysis reveals respiratory environmental traits involved in COVID-19 severity following SARS-CoV-2 infection. Nat Commun 2022, 13:4597.

58. Robinson O, Chadeau Hyam M, Karaman I, Climaco Pinto R, Ala-Korpela M, Handakas E, Fiorito G, Gao H, Heard A, Jarvelin MR, et al: Determinants of accelerated metabolomic and epigenetic aging in a UK cohort. Aging Cell 2020, 19:e13149.

59. Zhang X, Hu Y, Aouizerat BE, Peng G, Marconi VC, Corley MJ, Hulgan T, Bryant KJ, Zhao H, Krystal JH, Justice AC, Xu K: Machine learning selected smoking-associated DNA methylation signatures that predict HIV prognosis and mortality. Clin Epigenetics 2018, 10:155.

60. Ventham NT, Kennedy NA, Adams AT, Kalla R, Heath S, O’Leary KR, Drummond H, consortium IB, consortium IC, Wilson DC, et al: Integrative epigenome-wide analysis demonstrates that DNA methylation may mediate genetic risk in inflammatory bowel disease. Nat Commun 2016, 7:13507.

61. Zannas AS, Jia M, Hafner K, Baumert J, Wiechmann T, Pape JC, Arloth J, Kodel M, Martinelli S, Roitman M, et al: Epigenetic upregulation of FKBP5 by aging and stress contributes to NF-kappaB-driven inflammation and cardiovascular risk. Proc Natl Acad Sci U S A 2019, 116:11370–11379.

62. Flanagan JM, Brook MN, Orr N, Tomczyk K, Coulson P, Fletcher O, Jones ME, Schoemaker MJ, Ashworth A, Swerdlow A, Brown R, Garcia-Closas M: Temporal stability and determinants of white blood cell DNA methylation in the breakthrough generations study. Cancer Epidemiol Biomarkers Prev 2015, 24:221–229.

63. Johansson A, Enroth S, Gyllensten U: Continuous Aging of the Human DNA Methylome Throughout the Human Lifespan. PLoS One 2013, 8:e67378.

64. You C, Wu S, Zheng SC, Zhu T, Jing H, Flagg K, Wang G, Jin L, Wang S, Teschendorff AE: A cell-type deconvolution meta-analysis of whole blood EWAS reveals lineage-specific smoking-associated DNA methylation changes. Nat Commun 2020, 11:4779.

65. Hannum G, Guinney J, Zhao L, Zhang L, Hughes G, Sadda S, Klotzle B, Bibikova M, Fan JB, Gao Y, et al: Genome-wide methylation profiles reveal quantitative views of human aging rates. Mol Cell 2013, 49:359–367.

66. Balduzzi S, Rücker G, Schwarzer G: How to perform a meta-analysis with R: a practical tutorial. Evidence Based Mental Health 2019, 22:153–160.

